# Amitraz Toxicity in Resistant Varroa Mites Can Be Increased by Inhibiting ABCB1 Transporters

**DOI:** 10.1101/2025.04.23.650251

**Authors:** Julia D Fine, Tina Truong, Michelle C Lucadello, Eliza M Litsey, Zachary Lamas, Frank Rinkevich, Sascha Nicklisch

## Abstract

As critical pollinators of agricultural crops, honey bees (*Apis mellifera*) play a vital role in food production. Therefore, it is imperative to investigate and develop new methods to control *Varroa destructor*, a devastating parasitic mite of honey bee colonies that weakens bees and spreads deadly diseases that lead to colony loss. Here, we adapted existing methods to investigate the role of ABCB1 transporters in mitigating the toxicity of amitraz, a widely used miticide approved for use in honey bee colonies, demonstrating that a pharmacological inhibitor can synergistically increase amitraz toxicity compared to the equivalent dose of amitraz alone. Evaluations performed on the mites used in one of the described experiments revealed a high proportion of the test subject mites (88.5%) possessed the amitraz resistant genotype, indicating that even in resistant mites, inhibiting ABCB1 transporters can increase amitraz efficacy. This promising finding may be useful in developing powerful synergists that can be used to increase the efficacy of new and existent miticides.

## Introduction

Pollination by managed honey bee colonies is critical to the production of numerous agricultural crops in the United States and globally (Calderone, 2012; Southwick and Southwick, 1992), but, due to unrelenting stress caused by myriad biotic and abiotic factors, many beekeepers report experiencing unsustainable colony losses in yearly surveys (Aurell et al., 2024; Beyer et al., 2018; Brown et al., 2018; Bruckner et al., 2023; Rennich et al., 2014; vanEngelsdorp et al., 2013, 2009, 2008; Zee et al., 2015). Although the causes of these losses vary, parasitism by *Varroa destructor* is widely recognized as the most harmful and ubiquitous stressor of honey bee colonies (Jack and Ellis, 2021; Traynor et al., 2020). *Varroa destructor*, or *Varroa* mites, originated in Asia as parasites of the eastern honey bee, *Apis cerana*. Although the exact year that *Varroa destructor* shifted hosts to the western honey bee, *Apis mellifera*, is not known, it is thought to have occurred sometime in the first half of the 19^th^ century (Rosenkranz et al., 2010), and the first report of *Varroa* mites in the United States was made in 1987 (Guzman and Rinderer, 1999). Since then, they have become the most devastating and economically significant pest of honey bee colonies in the country (McMenamin and Genersch, 2015; Traynor et al., 2020, 2016).

*Varroa* mite reproduction occurs inside of honey bee brood cells, which a foundress mite enters just prior to capping (Boot et al., 1994; Ifantidis, 1983). Inside, the foundress mite and her offspring feed on the developing honey bee until it ecloses or is removed from the cell by workers (though the latter does not typically occur frequently enough in most *Apis mellifera* colonies to control mite populations) (Boecking and Spivak, 1999). Prior to the reproductive phase in the pupal cells, female adult *Varroa* mites feed on adult workers and drones (Ifantidis, 1983; Jack and Ellis, 2021). During this nonreproductive phase, *Varroa* mites can move from one colony to another as their honey bee hosts enter new colonies due to drift or robbing (Frey et al., 2011). In this way, *Varroa* infestations spread quickly and affect all colonies to varying degrees in a given area (Nolan and Delaplane, 2016). *Varroa* mites can harm honey bees by feeding on their tissues (Han et al., 2024; Ramsey, 2021) and by vectoring deadly viruses such as Deformed Wing Virus, Israeli Acute Paralysis Virus, and others (Gisder et al., 2009; Martin et al., 2012). This results in weakened and virus infected worker bees at the individual level, and population declines and losses at the colony level.

Despite the considerable damage caused by *Varroa* mites, beekeepers have limited tools available to control them. Cultural practices such as inducing brood breaks to prevent mites from reproducing (Gregorc et al., 2017; Lodesani et al., 2014) and removing drone comb from colonies can be somewhat effective (Calis et al., 1999; Wantuch and Tarpy, 2009), but these are not always feasible on a commercial scale and work best when combined with other control methods (Gregorc et al., 2017; Jack et al., 2020). Likewise, stocks of honey bees exhibiting resistance to *Varroa* are available (Bubnič et al., 2021; Danka et al., 2016; Rinderer et al., 2000; Traynor et al., 2020), but beekeepers often must select stocks due to other desirable traits such as the rate of colony build- up and temperament (Brother and Adam, 2000). By far, the most widely used methods to control *Varroa* mites are chemical (Haber et al., 2019; Jack and Ellis, 2021).

Relative to other agricultural markets, there are few pesticide products registered for use in honey bee colonies to control *Varroa* mites (Rosenkranz et al., 2010). Currently, one of the most widely used miticides for controlling *Varroa* is amitraz, a synthetic formamidine acaricide that mimics the activity of octopamine and tyramine (Casida and Durkin, 2013; Guo et al., 2021; R.m and A.e, 1982). Amitraz is typically applied inside colonies as the formulated and EPA approved products, Apivar®, via a slow-release strip and is thought to act via contact exposure as bees walk over the strip and spread amitraz throughout the colony or Amiflex® which is a fast release gel applied to top bars that provides rapid knockdown via contact exposure. Amitraz is preferred because of its high efficacy against *Varroa* mites and low toxicity to honey bees, but concerns surrounding its use have mounted in recent years due to the evolution of resistance in *Varroa* mite populations (Maggi et al., 2010; Rinkevich, 2020). This resistance is caused by a mutation in the ß2-octopamine receptor (Hernández-Rodríguez et al., 2025; Rinkevich et al., 2023). While a target-site resistance mutation affecting receptor selectivity can profoundly influence the toxicity of amitraz alone, the use of agricultural synergists targeting the biological factors underpinning metabolic resistance may offer promising solutions (Correy et al., 2019; Snoeck et al., 2017). It is possible that a synergist capable of influencing cellular uptake and accumulation of amitraz by inhibiting efflux transporters could help to overcome this resistance.

ATP-binding cassette (ABC) transporters involved in multidrug/xenobiotic resistance (MDR/MXR) are ubiquitously a first line of defense against the cellular uptake of a wide range of pesticide, drugs, and other toxic chemicals or metabolites(Galetin et al., 2024; Sauna et al., 2001). Through immediate recognition and eflux of substrate chemicals, MDR/MXR-type ABC transporters offer a rapid, energy-efficient, and versatile cellular defense response against chemical insults (Döring and Petzinger, 2014). While homologs of these eflux transporters have been identified across numerous arthropod taxa, repeatedly playing a major role in detoxification of broad-spectrum insecticides (Denecke et al., 2021; Dermauw and Van Leeuwen, 2014; Merzendorfer, 2014), the mechanisms governing pesticide recognition and cellular entry, orchestrated by the ABC gatekeeper proteins, have yet to be fully elucidated in *Varroa* and other pest organisms. If miticides are recognized as substrates for these transporters, their inhibition could increase cellular miticide accumulation, thereby sensitizing mites to their toxic effects. This process of chemosensitization has been described decades ago in aquatic species and has since been observed in other vertebrates and invertebrates (Epel et al., 2008; Kurelec, 1997; Nicklisch and Hamdoun, 2020).

One of the most ubiquitous and best characterized arthropod ABC transporters is MDR49, aka P-glycoprotein or ABCB1 (Gott et al., 2017). Natural mutation and overexpression of this protein are known to be critical to insecticide detoxification in mosquitos, tobacco budworm, and fruit flies (Denecke et al., 2022; Epis et al., 2014; Gellatly et al., 2015; Lanning et al., 1996b, 1996a; Seong et al., 2016; Seong and Pittendrigh, 2020). As a corollary, genetic knockdowns, multiple pesticide co-exposures, and pharmacological inhibition of these transporters can sensitize both pests and honey bees towards toxic chemical accumulation (Buss et al., 2002; Buss and Callaghan, 2008; Guseman et al., 2016; Hawthorne and Dively, 2011; Merzendorfer, 2014; Porretta et al., 2008). For instance, in fruit flies (*Drosophila melanogaster*), RNAi knockdown of the two P-glycoproteins MDR50 and MDR65 increased sensitivity towards DDT and ivermectin (Kim et al., 2018). Similarly, in the parasitic tick *Rhipicephalus microplus*, sensitivity towards ivermectin, abamectin, moxidectin and chlorpyriphos was enhanced by pharmacological inhibition of ABCB-type transporters using cyclosporine A (Pohl et al., 2014, 2012, 2011). In the red flour beetle (*Tribolium castaneum*), treatment with verapamil, an ABCB1 inhibitor, likewise increased susceptibility towards diflubenzuron (Rösner and Merzendorfer, 2020).

Here, we adapt existing methods to evaluate miticide toxicity and use them to investigate the effects of amitraz and the ABCB1 transporter inhibitor, verapamil, on *Varroa* mites both alone and in combination to elucidate the role of ABCB1 transporters in mitigating the toxicity of amitraz in *Varroa*. Additionally, we follow-up our experiments by quantifying the presence of the amitraz resistance genotype in the mites used for our late-season bioassays to better understand how ABC transporters may influence amitraz toxicity in resistant populations.

## Methods

### Mite Collection

Mites were obtained by powdered sugar dusting of colonies (Fakhimzadeh, 2001). Briefly, a third of a cup of powdered sugar per box was sprinkled through sieve over the top bars of colonies located at one of two apiaries located within five miles of the Pollinator Health Laboratory in Davis, CA, herein referred to as Apiaries A and B. The colonies were closed for twenty minutes, and the adult nonreproductive mites that detached from the adult bees were collected from the bottom boards of the colonies. Mites were maintained in the laboratory using methods adapted from a previously published protocol (Egekwu et al., 2018). The mites were taken directly to the Pollinator Health Laboratory, gently cleaned using damp cotton swabs, and groups of five mites were placed onto white or pink-eyed pupae obtained from colonies in the same apiary. Pupae were placed in size 00 gel caps and placed in a desiccator housed inside an incubator set to 34.5°C. Relative humidity was maintained at 75% inside the desiccator using a saturated aqueous solution of NaCl in a tray placed on the bottom. The mites remained in the desiccator for up to 3 hours until the treatments were applied.

### Amitraz and Verapamil Exposure

Mites were exposed to treatments using a microinjector to administer precise amounts of the treatment solutions to the mites’ dorsal cuticle using a method was adapted from Bahreini et al. (Bahreini et al., 2020) (Fig.1). Before administering the treatments, mites were placed in half of a 100 mm petri dish lined in wax foundation as mites were observed to be less mobile when the wax lining was used. Using the UMP3 Micro2T microinjector with a 33G beveled NanoFil needle (WPI Inc, Sarasota, FL), the treatment solution was applied to the dorsal cuticle of the mites at the volume specified below without permitting the liquid to contact the mouth, palps, or chelicerae. After the treatment was applied, mites were placed onto pupae in gel caps in groups of 5. Pupae were once again placed in a desiccator inside an incubator set at the previously mentioned conditions, and survival of the mites was checked at 24-hour intervals for up to 48 hours. Mite mortality was ascertained by a lack of movement observed after gently prodding the mite with a toothpick.

**Figure 1:**
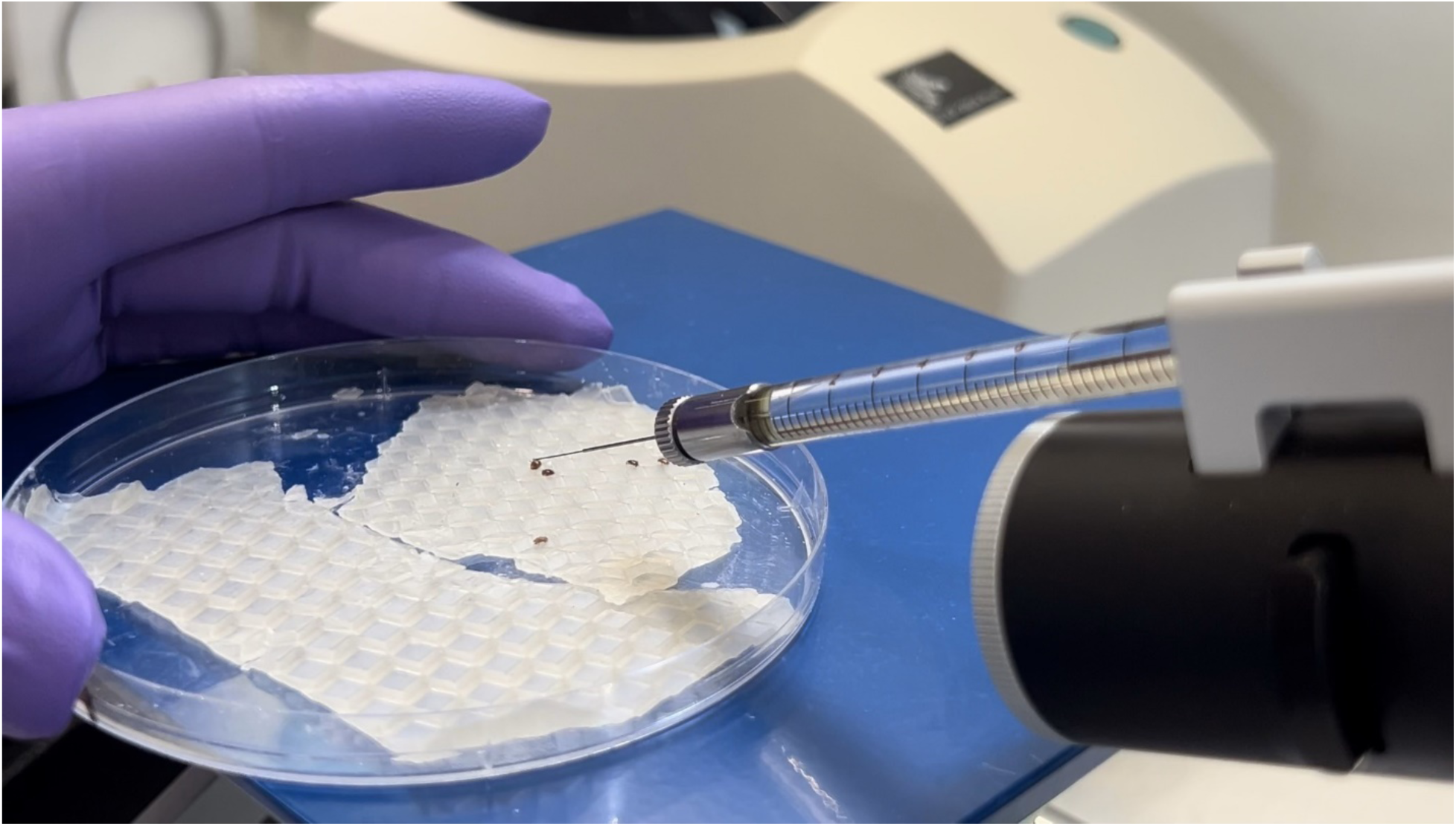
*Varroa* mites were placed on a petri dish lined in wax foundation, and treatments were applied to the dorsal cuticle of the mites using a microinjector.

### Treatments

#### Experiment 1

An initial range finding experiment was conducted on August 7, 2024 examining the effects of two doses of amitraz (0.0117 ng and 0.0234 ng/mite), two doses of verapamil (0.197 ng and 0.394 ng/mite). The dose of amitraz was based on the results of a previous study (Bahreini et al., 2020), which found, using similar methods, that exposing mites to 0.0117 ng of amitraz per mite resulted in 37.5% mortality in the tested population. The two doses of verapamil were obtained by multiplying the molar equivalents of the amitraz doses by ten. Additional treatments consisted of combinations of amitraz and verapamil (0.0117 ng amitraz + 0.197 ng verapamil per mite and 0.0234 ng amitraz + 0.394 ng verapamil per mite) and a solvent control. Each combination of amitraz and verapamil was administered in a single solvent application. All treatments were administered in 150 nL of acetone, and 15 mites were used per treatment group. If a mite died following the treatment before it could be placed onto a pupa it was excluded from the experiment, as we could not discern the cause of death from potential mishandling. This resulted in as few as 13 mites in certain treatment groups. Mites were obtained from two colonies located in Apiary A. Mite survival was assessed 24 hours after treatment application.

#### Experiment 2

Based on the results of Experiment 1, another miticide exposure experiment was conducted on August 14, 2024. This time, the amitraz dose was increased 4-fold relative to the low amitraz dose used in Experiment 1, resulting in a dose of 0.0468 ng per mite (amitraz-0.0468 ng). Based on the results of Experiment1, which demonstrated that 0.197 ng of verapamil alone did not cause mite mortality, a dose of 0.197 ng verapamil (verapamil) was used in Experiment 2. Once again, a combination of amitraz and verapamil administered in a single solvent application was also examined (0.0468 ng amitraz + verapamil) along with a solvent control (control). All treatments were administered in 150 nL of acetone, and 10 mites were used per treatment group. No mites were observed to be dead following the application of treatments before they could be placed on pupae. The experiment was replicated on August 19, 21, and 28. On August 19, 20 mites per treatment group were used, but due to mortality immediately after treatment, as few as 19 mites were used in certain treatment groups. On August 21 and 28, no mites were observed to be dead following the application of treatments before they could be placed on pupae, and 20 mites were used per treatment group on both dates. For all replicates of this experiment, mites were obtained from the same two colonies in Apiary A as those used in Experiment 1, and mite survival was assessed 24 hours after treatment application.

#### Experiment 3

On October 10, an experiment was performed to evaluate the toxicity of amitraz when combined with verapamil later in the season. Due to fluctuations in mite abundance in Apiary A, mites collected for this work from 2 colonies in Apiary B. Treatment groups consisted of 0.197 ng verapamil per mite (verapamil), 0.0468 ng amitraz per mite (0.0468 ng amitraz), and a group co-exposed to 0.197 ng verapamil and 0.0468 ng amitraz per mite administered in a single solvent application (0.0468 ng amitraz + verapamil). This time, treatment was administered in 300 nL of acetone, and to explore the potential effects of the increased solvent quantity on mite survival, we included two acetone control treatments: one exposed to 150 nL (150 nL control) and one exposed to 300 nL (300 nL control). The increased solvent volume was investigated as a means of alleviating the difficulty of administering a smaller volume of acetone to living *Varroa* mites before it evaporated. Up to 15 mites per treatment group were used, but due to mortality immediately after treatment, as few as 13 mites were used in certain treatment groups. Mite survival was assessed 24 hours after treatment application.

#### Experiment 4

On October 22, another experiment was performed on mites collected from the same 2 colonies in Apiary B, this time assessing the effects of an increased dose of amitraz in combination with verapamil. Treatment groups consisted of 0.197 ng verapamil per mite (verapamil), 0.374 ng amitraz per mite (0.374 ng amitraz), a group co-exposed to 0.197 ng verapamil and 0.374 ng amitraz per mite (0.374 ng amitraz + verapamil) administered in a single solvent application, and a solvent control (control). All treatments were administered in 300 nL acetone. Up to 20 mites per treatment group were used, but due to mortality immediately after treatment, as few as 18 mites were used in certain treatment groups.

This experiment was replicated on October 30, using mites collected from the same colonies exposed to the same treatments. Up to 17 mites per treatment group were used, but due to mortality immediately after treatment, as few as 16 mites were used in certain treatment groups. For both replicates, mite survival was assessed 24 and 48 hours after treatment application. At the conclusion of the experiment, both live and dead mites from Experiment 4 were collected and shipped to the USDA Honey Bee Breeding, Genetics, and Physiology Research Lab (HBBGPRL) in Baton Rouge, LA for amitraz resistance genotyping.

### Amitraz Resistance Genotyping

Genomic DNA was extracted from individual *Varroa* in a 2 mL tube with 2.8 mm ceramic beads with a Bead Ruptor Elite Mill (OMNI-International) for 2 cycles of 4 m/s for 5 s with a 5 s rest between cycles using water as a solvent. After homogenization, samples were centrifuged at 1000 x g for 2 min and the supernatant was used in a TaqMan assay to detect the Y215H mutation in the β2 octopamine receptor that is associated with amitraz resistance in *Varroa* (Hernández-Rodríguez et al., 2025; Rinkevich et al., 2023). All 134 of the *Varroa* sent to the HBBGPRL were genotyped. Statistical Analyses

To increase statistical power, survivorship was pooled across all replicates in an experiment. Normality and homogeneity of variances were assessed using Shapiro Wilk and Bartlett tests respectively. Based on the results of these, a non-parametric approach was used for all survivorship data. Differences in the proportions of mites surviving in each treatment group was assessed using Kruskal-Wallis rank sum tests (KW), and post hoc Dunn’s tests (DT) with Benjamini-Hochberg adjustments for multiple comparisons. Figures were created and statistical analyses were performed using R version 4.4.1. All data has been made available in supplemental data file S1.

## Results

No differences were observed in the proportion of mites surviving 24 hours after exposure to any treatments in Experiment 1 (KW, ꭓ^2^=8.27, df=5, p=0.142), and based on this result, the lowest dose of amitraz (0.0117 ng/mite) was increased by a factor of 4 in Experiment 2. A higher percentage of mites survived for 24 hours following exposure to the lower dose of verapamil (0.197 ng/mite – 86.7%) compared to the higher dose (0.394 ng/mite – 71.4%), though the difference in survival was not statistically significant. To allow for the largest possible synergism with verapamil while maintaining high survival rates in mites exposed to verapamil alone, the lower dose was used for the remaining experiments.

A significant difference in the proportion of mites surviving 24 hours after exposure to the treatments in Experiment 2 was detected (KW, ꭓ^2^=10.9, df=3, p=0.0124), and post hoc testing revealed that significantly fewer mites survived after exposure to 0.0468 ng amitraz together with 0.197 ng verapamil/mite compared to mites exposed to the solvent control or verapamil alone (DT, Z=-3.06, p=0.0132 and Z=-2.60, 0.0279 respectively, Fig. 2).

**Figure 2:**
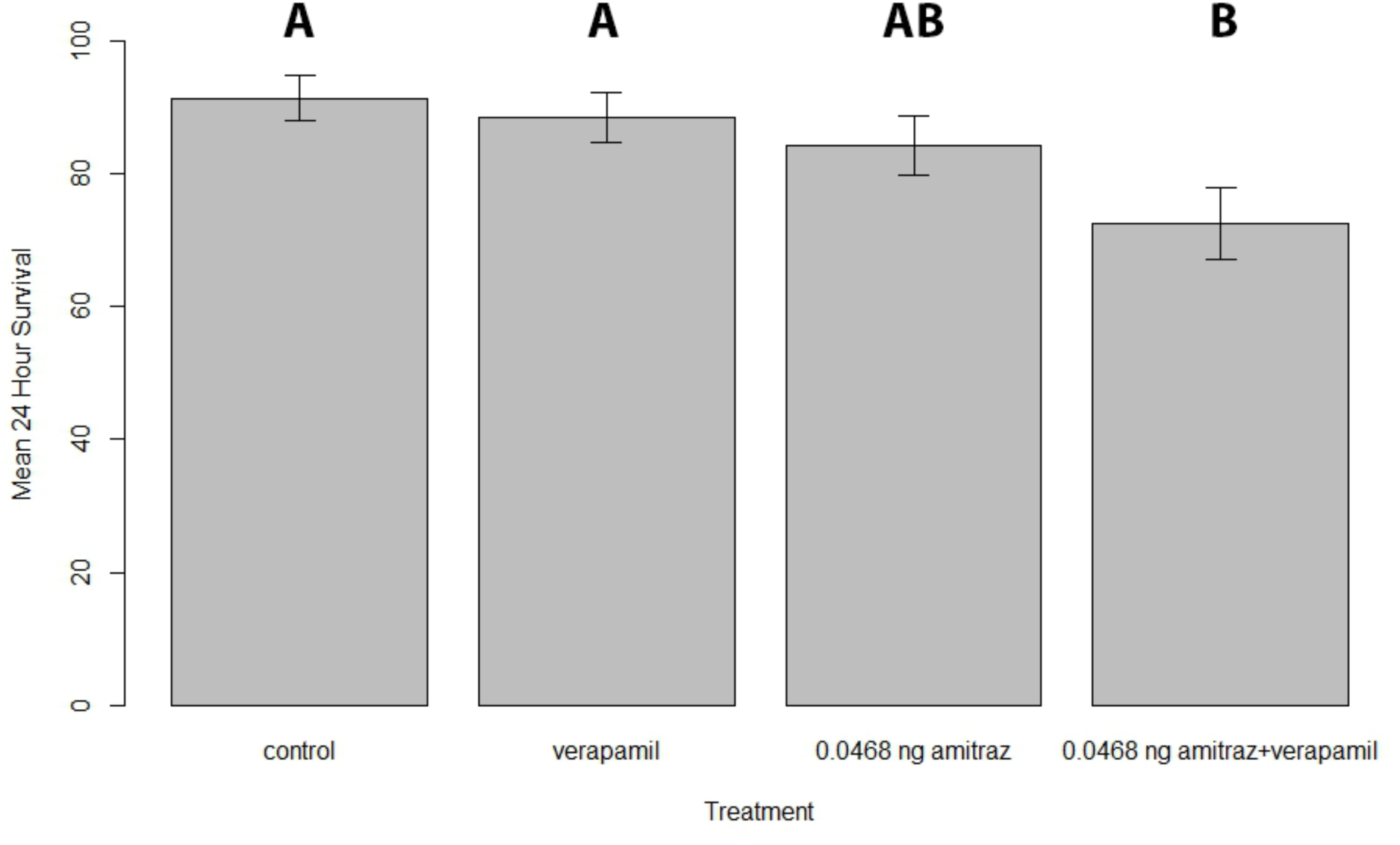
Mean percent of surviving mites 24 hours after treatment in Experiment 2 with a solvent control, amitraz and verapamil along and in combination (KW, ꭓ^2^=10.9, df=3, p=0.0124, significant differences correspond to nonoverlapping letter groups).

In Experiment 3, no significant differences were detected between treatment groups (KW, ꭓ^2^=5.08, df=4, p=0.280), suggesting that using 300 nL of solvent would not affect the survival of mites.

However, no effect of amitraz (at 1 and 2x the previously tested dose) alone or in combination with verapamil was observed, suggesting that a higher dose may be needed to influence the survival of the mites taken from colonies in Apiary B in October. Therefore, the highest dose of amitraz tested in Experiment 3 was increased by a factor of 8 for Experiment 4.

After 24 hours, a significant difference in the proportion of mites was detected among treatment groups in Experiment 4 (KW, ꭓ^2^=9.24, df=3, p=0.0263), but post hoc testing revealed that the only difference was that a lower proportion of mites co-exposed to 0.374 ng amitraz and 0.197 ng verapamil survived relative to mites exposed to verapamil alone (DT, Z=-2.42, p=0.0471, Fig. 3). However, after 48 hours, significant differences among treatment groups were again detected (KW, ꭓ^2^=38.05, df=3, p≤0.001). At this timepoint, post hoc tests revealed that a significantly lower proportion of mites co-exposed to 0.374 ng amitraz and 0.197 ng verapamil survived relative to all other treatment groups (DT, Control: Z=-5.01, p≤0.001; 0.197 ng verapamil: Z=-5.48, p≤0.001; 0.374 ng amitraz: -2.71, p=0.010, Fig. 3) and a significantly lower proportion of mites exposed to 0.374 ng amitraz survived relative to control and 0.197 ng verapamil/mite (Z=-2.36, p=0.022 and Z=-2.75, p=0.012 respectively).

**Figure 3.**
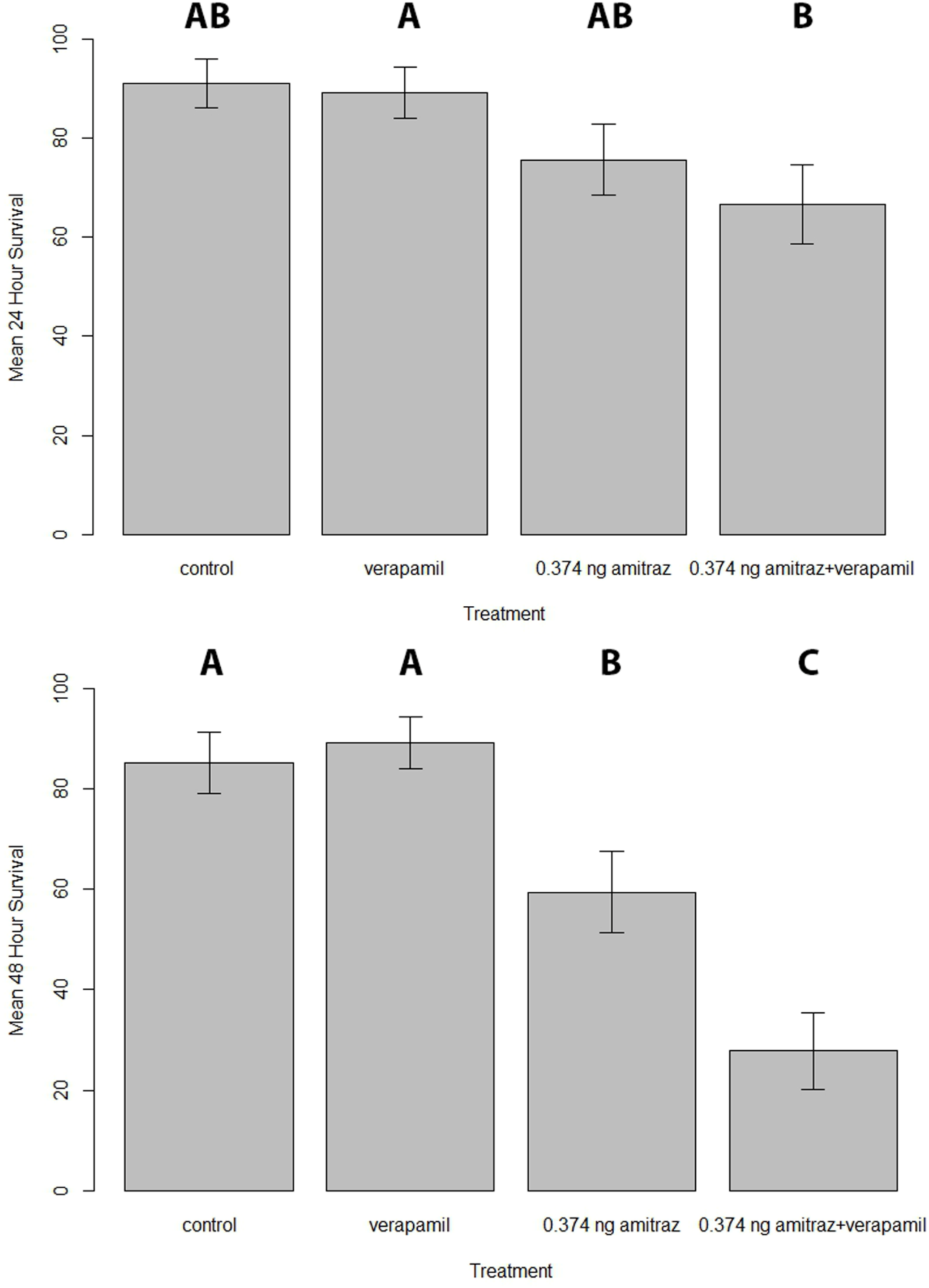
Top: Mean percent of surviving mites 24 hours after treatment in Experiment 4 with a solvent control, amitraz and verapamil along and in combination (KW, ꭓ^2^=9.24, df=3, p=0.0263, significant differences correspond to nonoverlapping letter groups). Bottom: Mean percent of surviving mites 48 hours after treatment in Experiment 4 with a solvent control, amitraz alone, verapamil alone, and a combination of amitraz and verapamil (KW, ꭓ^2^=38.05, df=3, p≤0.001, significant differences correspond to nonoverlapping letter groups).

The overall percentage of homozygous amitraz resistant genotypes for the *Varroa* tested was 88.5% and the percentages were not significantly different among treatment groups (Likelihood Ratio, χ2=10.408, p=0.4055, n=134, Fig. 4).

**Figure 4.**
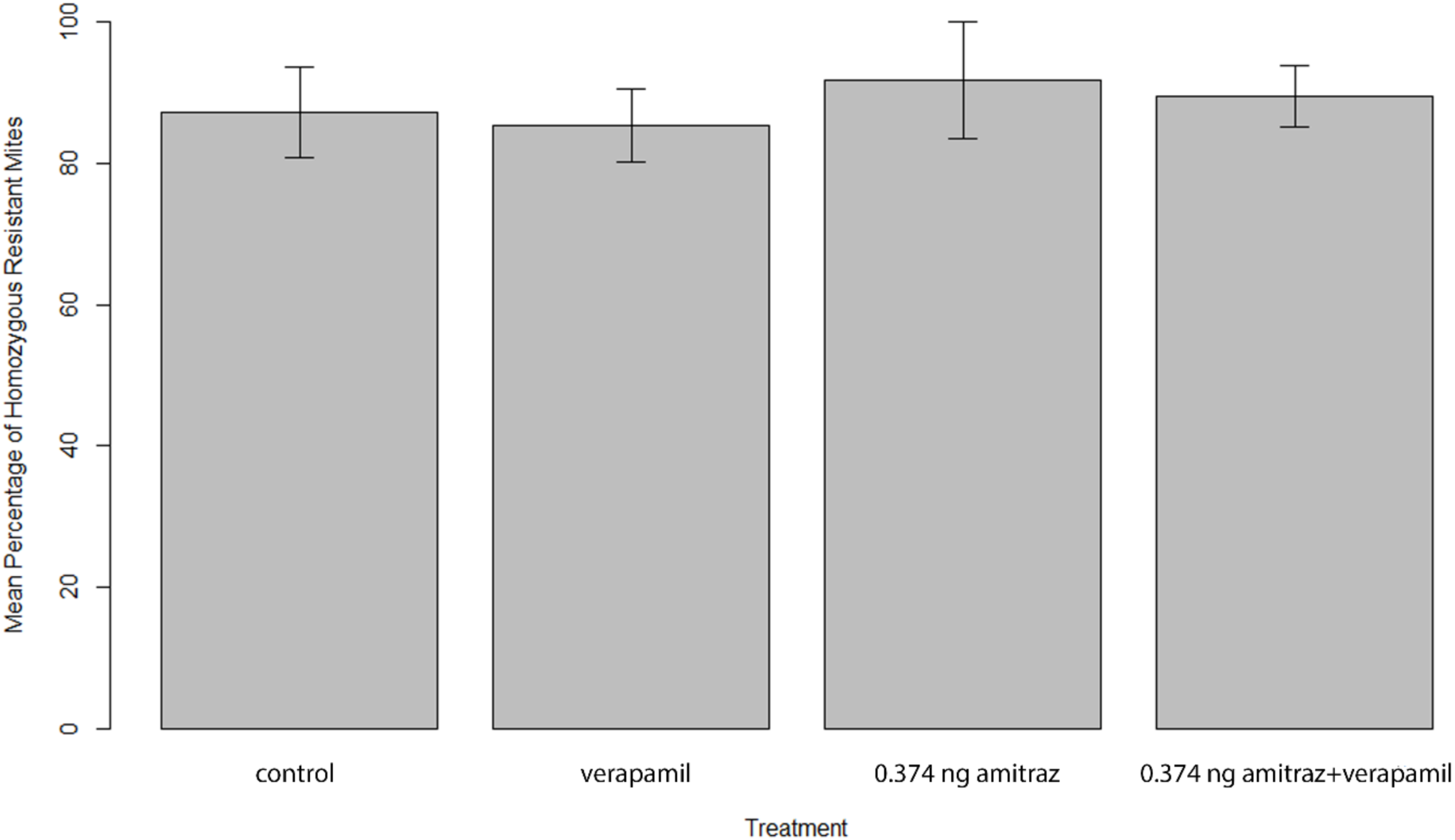
Top: Mean percentage of mites (± SE) possessing the resistant genotype by treatment. No difference in the percentage of resistant mites was found between treatment groups (Likelihood Ratio, χ2=10.408, p=0.4055, n=134).

## Discussion

In drug transporter research, pharmacological inhibition is often used as a control to identify and validate substrates for a transporter (Galetin et al., 2024; Nicklisch and Hamdoun, 2020). One of the best characterized and potent inhibitors of ABCB1 transporters, verapamil, has been successfully shown to synergize tick mortality with different miticides and insecticides that are known ABCB1 substrates (Shakya et al., 2022). Indeed, in this study on *Varroa* co-exposure to amitraz and verapamil, we confirmed their synergistic mortality, indicating that amitraz is a substate for *Varroa* ABCB1 which can be pharmacologically inhibited by verapamil. In our initial experiment, we did not observe significant mortality in mites treated with a dose of amitraz found to be toxic in a previous study, and the addition of verapamil did not increase mortality. When the experiment was repeated with a four-fold higher dose of amitraz, significant mortality was only observed when mites were co-exposed to verapamil. Later in the season, when *Varroa* were sourced from different colonies, a higher dose equivalent to roughly 32 times the initial dose of 0.0117 ng/mite had to be used to elicit any effect, however, in this experiment, dramatic effects were not observed until 48 hours after treatment as opposed to 24 hours. This is consistent with the very high levels of amitraz resistance genotypes observed in this study. Therefore, the amitraz resistance level from bioassays or genotyping should be considered in research using amitraz on *Varroa* in order to explain discrepancies in expected results.

Overall, the results reported here confirm that inhibiting *Varroa* ABCB1 transporters can increase the toxicity of amitraz, likely by decreasing cellular efflux of amitraz, but the increasing dose of amitraz needed to elicit an effect suggests that the mites in some or all of our experiments may have exhibited resistance to amitraz. Indeed, this was confirmed for the mites used in Experiment 4, which required a dose that was roughly 28 times a previously reported LD50 for amitraz to elicit an effect (Bahreini et al., 2020). A high proportion of these mites (88.5%) were homozygous for the resistant genotype. However, even in these resistant mites, mortality was increased by roughly 50% when mites were co-exposed to amitraz and verapamil relative to amitraz alone after 48 hours. Increased mortality was also observed within 24 hours in the co-exposed group, though only relative to the verapamil control group. This indicates that while resistant mites are still less susceptible to amitraz in the presence of an ABCB1 transporter inhibitor, the inhibitor does increase amitraz toxicity in this less susceptible population. This synergism with verapamil (or another suitable inhibitor) may extend the utility of amitraz for *Varroa* control and reduce the probability of the development of high levels of amitraz resistance.

Despite the importance of ABCB1-type efflux transporters, as demonstrated here and in other studies performed on arthropods (Buss and Callaghan, 2008; Denecke et al., 2022; Guseman et al., 2016; Kim et al., 2018; Lanning et al., 1996b; Pohl et al., 2012; Porretta et al., 2008; Rösner and Merzendorfer, 2020), the majority of research on pesticide detoxification mechanisms in numerous species including *Varroa* has been the cytochrome P450 enzyme complex, which is broadly involved in xenobiotic transformation and detoxification (De Rouck et al., 2023; Van Leeuwen and Dermauw, 2016; Vlogiannitis et al., 2021; Ye et al., 2022). Synergism bioassays performed on ticks and mites, where a miticide is combined with a specific inhibitor of Phase I and Phase II enzymes, show an increase in sensitivity towards the miticide in the presence of the inhibitors and indicate that metabolic detoxification mechanisms could be involved in resistance in some strains (Bernard and Philogène, 1993; Guerrero et al., 2012; Li et al., 2004, 2003; Lima et al., 2013). Because of this activity, inhibitors of cytochrome P450 enzymes have been used in agriculture for almost a century to synergize the effects of P450 detoxified pesticides (Bernard and Philogène, 1993). Given the pervasive role of ABCB1 in mediating toxic effects of pesticides in arthropods as well as their role in *Varroa* detoxification established by this work, exploiting their role in *Varroa* chemosensitization for pest control is a logical step. This process of chemo-sensitization, described decades ago, has already been successfully applied in pharmacology to treat human cancer cells and reverse multidrug resistance (Choi, 2005; Giacomini et al., 2010; Giacomini and Huang, 2013; Robey et al., 2018; Wu et al., 2008). However, because of the close association between *Varroa* mites and honey bees, care must be taken to develop and identify inhibitors of ABCB1 transporters that do not affect honey bee transporters.

In honey bees, ABCB1 transporters have been found to be involved in mitigating pesticide toxicity (Guseman et al., 2016; Hawthorne and Dively, 2011). ABCB1 shows high evolutionary conservation of structure and function across species from bacteria to humans (Bosch and Croop, 1998). This functional conservation includes promiscuous ligand-binding sites for substrate and inhibitor recognition (Alam et al., 2019; Aller et al., 2009; Le et al., 2020). Because of this conservation, using a universal pharmacological ABCB1 inhibitor like verapamil inside of a colony environment to control *Varroa* mites could also chemosensitize honey bees toward toxic chemical accumulation. However, species-specific differences in ABCB1 transporter function exist and have only been recently studied in detail in the context of drug development (Azimi et al., 2023) and environmental toxicology (Nicklisch et al., 2021).

A more elegant approach to rapidly identify species-specific inhibitors of ABCB1 transporters could be the use of computer-aided drug discovery tools (Moesgaard et al., 2023; Namasivayam et al., 2021) By combining homology modeling and molecular docking analysis, a myriad of compounds could be screened to identify candidate chemicals that inhibit *Varroa* ABCB1 transporters without affecting honey bee transporters (Guéniche et al., 2024; Mora Lagares et al., 2020; Mora Lagares and Novič, 2022; Vilar et al., 2019). Similar approaches have already been successfully applied to identify specific inhibitors of pest target molecules, including essential enzymes and transporters (Chen et al., 2024). After *in vivo* validation, such chemicals could be used to synergize both existent miticides like amitraz and others that may be identified in the future to be used with a novel synergist. In the case of amitraz, such an approach would benefit beekeepers and may slow the evolution of resistance. By increasing the efficacy of an initial amitraz treatment, the need for additional treatments is decreased, lowering the selection pressure on the mite population and decreasing the economic burden on beekeepers. Similar approaches have been successfully implemented with established integrated resistance management (IRM) strategies that apply saturation with synergists and high dose/refuge (HDR) approaches to prevent rapid resistance development (Lester, 2023; Sudo et al., 2018).

Ultimately, care must be taken when developing any new pesticidal product, particularly one that must be used inside honey bee colonies. However, given the importance of honey bee pollination for food security (Calderone, 2012; Southwick and Southwick, 1992) and the increasing challenge of miticide resistance in *Varroa*, investigating and exploiting the susceptibilities of *Varroa* should be considered a priority for researchers. This work suggests that chemosensitization via inhibition of ABCB1 transporters shows great promise and may lead to new advancements for *Varroa* control, including enhanced efficacy of the limited arsenal of approved miticides and potentially reduced treatment frequency.

## Acknowledgements

Mention of trade names or commercial products in this publication is solely for the purpose of providing specific information and does not imply recommendation or endorsement by the U.S. Department of Agriculture. USDA is an equal opportunity provider and employer.

## Data Availability

All data has been made available in supplemental data file S1.

---

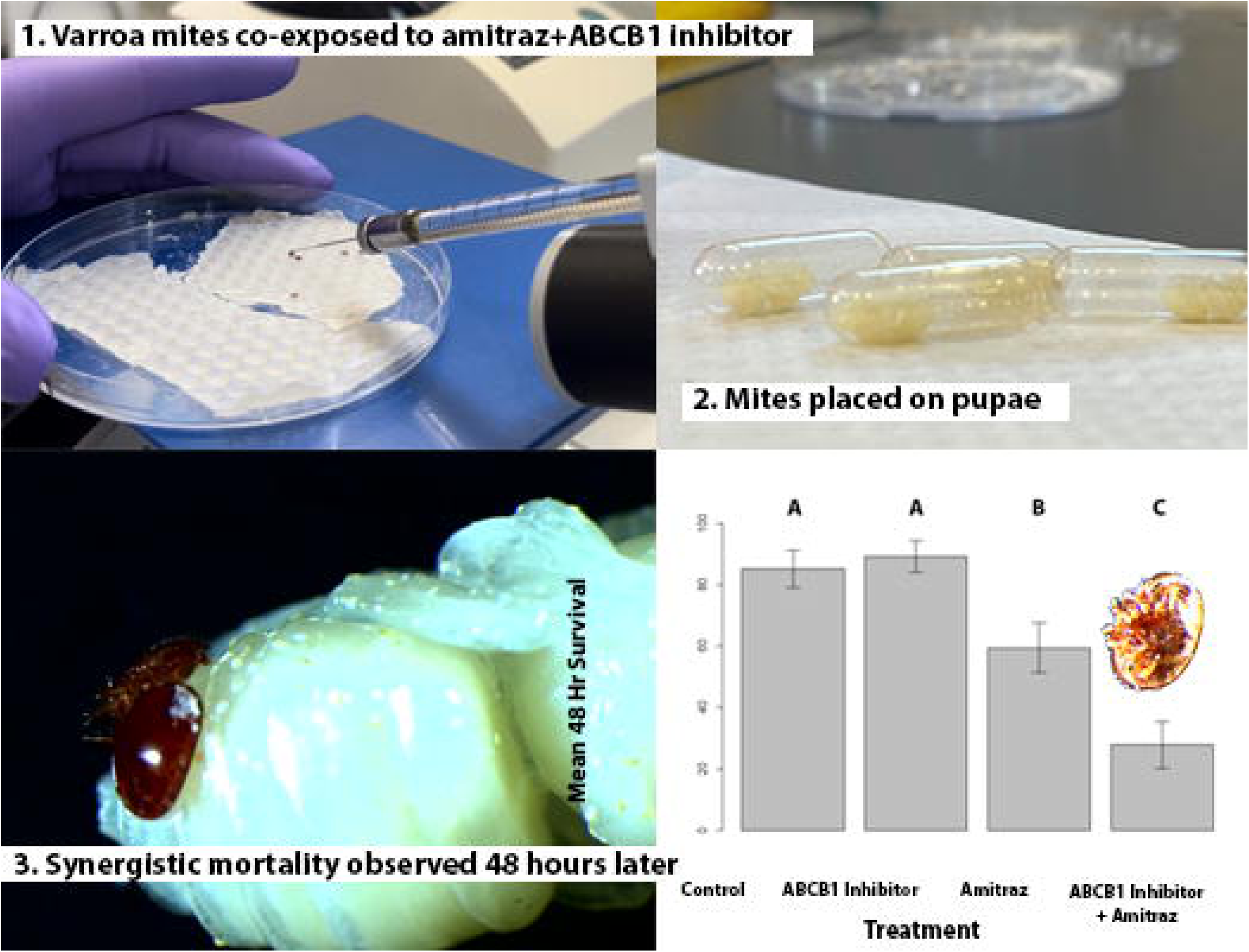

